# *Streptococcus pneumoniae* evades host cell phagocytosis and limits host mortality through its cell wall anchoring protein PfbA

**DOI:** 10.1101/599001

**Authors:** Masaya Yamaguchi, Yujiro Hirose, Moe Takemura, Masayuki Ono, Tomoko Sumitomo, Masanobu Nakata, Yutaka Terao, Shigetada Kawabata

## Abstract

*Streptococcus pneumoniae* is a Gram-positive bacterium belonging to the oral streptococcus species, mitis group. This pathogen is a leading cause of community-acquired pneumonia, which often evades host immunity and causes systemic diseases, such as sepsis and meningitis. Previously, we reported that PfbA is a β-helical cell surface protein contributing to pneumococcal adhesion to and invasion of human epithelial cells in addition to its survival in blood. In the present study, we investigated the role of PfbA in pneumococcal pathogenesis. Phylogenetic analysis indicated that the *pfbA* gene is specific to *S. pneumoniae* within the mitis group. Our *in vitro* assays showed that PfbA inhibits neutrophil phagocytosis, leading to pneumococcal survival. We found that PfbA activates NF-κB through TLR2, but not TLR4. In addition, TLR2/4 inhibitor peptide treatment of neutrophils enhanced the survival of the *S. pneumoniae* Δ*pfbA* strain as compared to a control peptide treatment, whereas the treatment did not affect survival of a wild-type strain. In a mouse pneumonia model, the host mortality and level of TNF-α in bronchoalveolar lavage fluid were comparable between wild-type and Δ*pfbA*-infected mice, while deletion of *pfbA* increased the bacterial burden in bronchoalveolar lavage fluid. In a mouse sepsis model, the Δ*pfbA* strain demonstrated significantly increased host mortality and TNF-α levels in plasma, but showed reduced bacterial burden in lung and liver. These results indicate that PfbA may contribute to the success of *S. pneumoniae* species by inhibiting host cell phagocytosis, excess inflammation, and mortality.

**Importance:** *Streptococcus pneumoniae* is often isolated from the nasopharynx of healthy children, but the bacterium is also a leading cause of pneumonia, meningitis, and sepsis. In this study, we focused on the role of a cell wall anchoring protein, PfbA, in the pathogenesis of *S. pneumoniae-*related disease. We found that PfbA is a pneumococcus-specific anti-phagocytic factor that functions as a TLR2 ligand, indicating that PfbA may represent a pneumococcal-specific therapeutic target. However, a mouse pneumonia model revealed that PfbA deficiency reduced the bacterial burden, but did not decrease host mortality. Furthermore, in a mouse sepsis model, PfbA deficiency increased host mortality. These results suggest that *S. pneumoniae* optimizes reproduction by regulating host mortality through PfbA; therefore, PfbA inhibition would not be an effective strategy for combatting pneumococcal infection. Our findings underscore the challenges involved in drug development for a bacterium harboring both commensal and pathogenic states.

## Introduction

*Streptococcus pneumoniae* is Gram-positive bacteria belonging to the mitis group that colonizes the human nasopharynx in approximately 20% of children without causing clinical symptoms (1-3). On the other hand, *S. pneumoniae* is also a leading cause of bacterial pneumonia, meningitis, and sepsis worldwide. The pathogen is estimated to be responsible for the deaths of approximately 1,190,000 people annually from lower respiratory infection (4). Following the introduction of pneumococcal conjugate vaccines, *S. pneumoniae* is still responsible for two thirds of all cases of meningitis (5). In addition, antibiotic selective pressure causes resistant pneumococcal clones to emerge and expand all over the world and the World Health Organization listed *S. pneumoniae* as one of antibiotic-resistant ‘priority pathogens’ (6). Centers for Disease Control and Prevention data from active bacterial core surveillance for 2009 to 2013 indicated that pneumococcal conjugate vaccines work as a useful tool against antibiotic resistance (7). However, these vaccines also generate selective pressure, and non-vaccine serotypes of *S. pneumoniae* are increasing worldwide (8, 9).

During the process of invasive infection, *S. pneumoniae* needs to evade host immunity and replicate in the host after colonization. In these steps, pneumococcal cell surface proteins work as adhesins and/or anti-phagocytic factors. There are two types of motifs for pneumococcal cell surface localization, a cell wall anchoring motif, LPXTG (10), and choline-binding repeats interacting with pneumococcal phosphorylcholine (11). Choline-binding proteins (CBPs) localize on the pneumococcal cell wall via the phosphorylcholine moiety of teichoic acids, while LPXTG-anchored proteins are covalently attached to the cell wall. Several LPXTG-anchored proteins and CBPs contribute to the adhesion to host epithelial cells through the interaction with host factors (10-13). Some pneumococcal cell surface proteins also contribute to bacterial survival by limiting complement deposition or inhibiting phagocytosis (11, 14-17). On the other hand, the host recognizes *S. pneumoniae* and regulates immune responses using pattern recognition receptors, including the Toll-like receptors (TLRs), nucleotide oligomerization domain-like receptors, and retinoic acid-inducible gene-I-like receptors (18). In addition, extracellular bacteria are recognized by TLR2 and TLR4 located on the host cell surface. TLR2 recognizes pneumococcal cell wall components and lipoproteins, while TLR4 senses a pore-forming toxin, pneumolysin (18, 19). Generally, both TLR2 and TLR4 agonists induce neutrophil activation and inhibit the apoptosis (20). However, in mouse influenza A virus and *S. pneumoniae* co-infection model, a TLR2 agonist decreased inflammation and reduced bacterial shedding and transmission (21). TLRs play important, but redundant, roles in the host defense and regulating inflammatory responses against pneumococcal infection. Appropriate immune responses contribute to pneumococcal clearance, while excessive inflammation can lead to serious tissue damage.

We previously reported that plasmin- and fibronectin-binding protein A (PfbA) plays a role in fibronectin-dependent adhesion to and invasion of epithelial cells, and that an *S. pneumoniae* PfbA-deficient mutant strain exhibited decreased survival in human blood (22, 23). PfbA is an LPXTG-anchored protein that features a right-handed parallel β-helix with a groove or cleft, formed by three parallel β-sheets and connecting loops (24, 25). Since the distribution and structural arrangement of the groove residues in the β-helix make it favorable for binding to carbohydrates, PfbA binds to D-galactose, D-mannose, D-glucosamine, D-galactosamine, *N*-acetylneuraminic acid, D-sucrose, and D-raffinose (26). PfbA also binds to human erythrocytes by interacting with *N*-acetylneuraminic acids on the cells (27).

In this study, we investigated the role of PfbA in pneumococcal pathogenesis. Phylogenetic analysis indicated that *pfbA* is specific to *S. pneumoniae* among the mitis group *Streptococcus*. Our *in vitro* analysis revealed that PfbA works as an anti-phagocytic factor and that the protein causes NF-κB activation via TLR2. In addition, Toll-interleukin 1 receptor adaptor protein (TIRAP) inhibition increased the survival rate of the *pfbA* mutant strain after incubation with neutrophils, while the wild-type (WT) strain was not affected. Mouse infection assays suggested that PfbA contributes to pneumococcal survival in at least some organs. However, in a mouse sepsis model, *pfbA* mutant strain-infected mice showed significantly higher mortality and TNF-α levels in blood. Our findings indicate that PfbA is a pneumococcus-specific anti-phagocytic factor and suppresses host excess inflammation.

## Materials and Methods

### Bacterial strains and construction of mutant strain

*Streptococcus pneumoniae* strains were cultured in Todd-Hewitt broth (BD Biosciences, San Jose, CA, USA) supplemented with 0.2% yeast extract THY medium, BD Biosciences) at 37°C. For selection and maintenance of mutants, spectinomycin (Fujifilm Wako Pure Chemical Corporation, Osaka, Japan) was added to the medium at 120 µg/mL. The *Escherichia coli* strain XL10-Gold (Agilent, Santa Clara, CA, USA) was used as a host for derivatives of plasmid pQE-30. All *E. coli* strains were cultured in Luria-Bertani (LB) broth supplemented with 100 µg/mL carbenicillin (Nacalai Tesque, Kyoto, Japan) at 37°C with agitation.

*S. pneumoniae* TIGR4 isogenic *pfbA* mutant strains were generated as previously described with minor modifications (22, 28, 29). Briefly, the upstream region of *pfbA*, an *aad9* cassette, the downstream region of *pfbA*, and pGEM-T Easy vector (Promega, Madison, WI, USA) were amplified by PrimeSTAR^®^ MAX DNA Polymerase (TaKaRa Bio, Shiga, Japan) using the specific primers listed in Supplementary Table 1. The DNA fragments were assembled using a GeneArt^®^ Seamless Cloning and Assembly Kit (Thermo Fisher Scientific, Waltham, MA, USA). The constructed plasmid was then transformed into *E. coli* XL-10 Gold, and the inserted DNA region was amplified by PCR. The products were used to construct mutant strains by double-crossover recombination with the synthesized competence-stimulating peptide-2. The mutation was confirmed by PCR amplification of genomic DNA isolated from the mutant strain.

### Cell culture

Human promyelocytic leukemia cells (HL-60, RCB0041) were purchased from RIKEN Cell Bank (Ibaraki, Japan). HL-60 cells were maintained in RPMI 1640 medium (Thermo Fisher Scientific) supplemented with 10% FBS, and were incubated at 37°C in 5% CO_2_. HL-60 cells were differentiated into neutrophil-like cells for 5 days in culture media containing 1.2% DMSO (30, 31). Cell differentiation was confirmed by nitro blue tetrazolium reduction assay (30).

Human TLR2/NF-κB/SEAP stably transfected HEK293 cells and human TLR4/MD-2/CD14/NF-κB/SEAP stably transfected HEK293 cells (Novus Biologicals, Centennial, CO, USA, currently sold by InvivoGen, San Diego, CA, USA) were maintained in DMEM with 4.5 g/L glucose, 10% FBS, 4 mM L-glutamine, 1 mM sodium pyruvate, 100 units/mL penicillin, 100 µg/mL streptomycin, 10 µg/mL blasticidin, and 500 µg/mL G418 and DMEM with 4.5 g/L glucose, 10% FBS, 4 mM L-glutamine, 1 mM sodium pyruvate, 100 units/mL penicillin, 100 µg/mL streptomycin, 10 µg/mL blasticidin, 2 µg/mL puromycin, 200 µg/mL zeocin, and 500 µg/mL G418, respectively. A secreted alkaline phosphatase reporter assay was performed according to the manufacturer’s instructions (Novus Biologicals).

### Phylogenetic analysis

Phylogenetic analysis was performed as described previously (17, 32, 33), with minor modifications. Briefly, homologues and orthologues of the *pfbA* gene were searched using tBLASTn (34). The sequences were aligned using Phylogears2 (35, 36) and MAFFT v.7.221 with an L-INS-i strategy (37), and ambiguously aligned regions were removed using Jalview (38, 39). The best-fitting codon evolutionary models for phylogenetic analyses were determined using Kakusan4 (40). Bayesian Markov chain Monte Carlo analyses were performed with MrBayes v.3.2.5 (41), and 4 × 10^6^ generations were sampled after confirming that the standard deviation of split frequencies was < 0.01. To validate phylogenetic inferences, maximum likelihood phylogenetic analyses were performed with RAxML v.8.1.20 (42). Phylogenetic trees were generated using FigTree v.1.4.2 (43) based on the calculated data.

### Human neutrophil and monocyte preparation

Human blood was obtained via venipuncture from healthy donors after obtaining informed consent. The protocol was approved by the institutional review boards of Osaka University Graduate School of Dentistry (H26-E43). Human neutrophils and monocytes were prepared using Polymorphprep (Alere Technologies AS, Oslo, Norway), according to the manufacturer’s instructions. Human blood was carefully layered on the Polymorphprep solution in centrifugation tubes, which were then centrifuged at 450 × *g* for 30 min in a swing-out rotor at 20°C. Monocyte and neutrophil fractions were transferred into tubes containing ACK buffer (0.15 M NH_4_Cl, 0.01 M KHCO_3_, 0.1 mM EDTA), then centrifuged, washed in phosphate-buffered saline, and resuspended in RPMI 1640 medium.

### Neutrophil bactericidal assays

The pneumococcal cells grown to the mid-log phase were resuspended in PBS. TIGR4 strains (3-11 × 10^3^ CFUs/well) with or without rPfbA (0, 10, or 100 nM) were combined with human neutrophils or neutrophil like-differentiated HL-60 cells (2 × 10^5^ cells/well), and R6 strains (1.4-2.0 × 10^2^ CFUs/well) were combined with human neutrophils (1 × 105 cells/well). The mixture was incubated at 37°C in 5% CO_2_ for 1, 2, and 3 h. Viable cell counts were determined by plating diluted samples onto TS blood agar. The growth index was calculated as the number of CFUs at the specified time point/number of CFUs in the initial inoculum. Bacterial phagocytosis was blocked by addition of cytochalasin D (20 µM), and pneumococcal killing was blocked by protease inhibitor cocktail set V (Merck, Darmstat, Germany; 500 µM AEBSF, 150 nM Aprotinin, 1 µM E-64, and 1 µM leupeptin hemisulfate, EDTA-free) at 1 h before incubation. To determine whether TLR2 and TLR4 signaling affect pneumococcal survival, 100 µM TIRAP (TLR2 and TLR4) inhibitor peptide or control peptide (Novus Biologicals) were added to neutrophils at 1 h before incubation.

### Time-lapse microscopic analysis

For time-lapse observations, isolated neutrophils were resuspended in RPMI 1640 at 1 × 10^6^ cells/mL. Next, 10 µL of *S. pneumoniae* R6 wild type or Δ*pfbA* strains (1 × 10^6^ CFUs) was added to 2 mL of the cells, and the mixture was incubated and observed at 37°C. Time-lapse images were captured using an Axio Observer Z1 microscope system (Carl Zeiss, Oberkochen, Germany).

### Flow cytometric analysis of phagocytes

Recombinant PfbA (rPfbA) or BSA was coated onto 0.5-µm-diameter fluorescent beads (FluoroSphere, Thermo Fisher Scientific), according to the manufacturer’s instructions. rPfbA was purified as previously described (22). Isolated neutrophils or monocytes were then resuspended in RPMI 1640 at 1.0 × 10^7^ cells/mL, after which 900 µL of RPMI 1640 containing 1 µL of rPfbA-, BSA-, or non-coated fluorescent beads was added to 100 µL of cells, and then the mixtures were rotated at 37°C for 1 h. The cells were washed twice and fixed with 2% glutaraldehyde-RPMI 1640 at 37°C for 1 h, then washed again three times and analyzed with a CyFlow flow cytometer (Sysmex, Hyogo, Japan) using FlowJo software ver. 8.3.2 (BD Biosciences, Franklin Lakes, NJ, USA).

### TLR2/4 SEAPorter assay

HEK cells expressing TLR2 or TLR4 were stimulated with *S. pneumoniae* and/or rPfbA for 16 h, according to the manufacturer’s instructions (Novus Biologicals). To avoid the effect of bacterial replication on this assay, *S. pneumoniae* were pasteurized by incubation at 56°C for 30 min. To perform the assay under the same condition, rPfbA was also incubated at 56°C for 30 min. Lipopolysaccharides from *Escherichia coli* O111:B4 (Sigma-Aldrich Japan Inc., Tokyo, Japan) for the TLR-4 cell line and Pam3CSK4 and Zymozan (Novus Biologicals) for the TLR-2 cell line were used as positive controls under the same conditions. Secreted alkaline phosphatase (SEAP) was analyzed using the SEAPorter Assay (Novus Biologicals) according to the manufacturer’s instructions. Quantitative data (ng/mL) were obtained using a standard curve for the SEAP protein.

### RNA extraction and miRNA array

We performed microRNA array analysis using neutrophil like-differentiated HL60 cells incubated with *S. pneumoniae* strains and/or 100 nM rPfbA for 1 h. We compared rPfbA-treated and non-treated cells, wild type and Δ*pfbA*-infected cells, and Δ*pfbA* with and without rPfbA-infected cells. In each cell sample, six replicates were pooled and total RNA including microRNA was isolated from the pooled cells by miRNeasy Mini Kit (Qiagen, Hilden, Germany). Approximately 1000 ng RNA was used for microarray analysis using Affymetrix GeneChip miRNA 4.0 arrays (Affymetrix, Santa Clara, CA, USA) through Filgen Inc. (Nagoya, Japan). Briefly, the quality of total RNA was assessed using a Bioanalyzer 2100 (Agilent). Hybridization was performed using a FlashTag Biotin HSR RNA Labeling Kit, GeneChip Hybridization Oven 645, and GeneChip Fluidics Station 450. The arrays were scanned by Affymetrix GeneChip Scanner 3000 7G. The GeneChip miRNA 4.0 arrays contain 30,424 total mature miRNA probe sets including 2,578 mature human miRNAs, 2,025 pre-miRNA human probes, and 1,196 Human snoRNA and scaRNA probe sets.

### Mouse infection assays

Mouse infection assays were performed as previously described (17, 33, 44, 45). For the lung infection model, CD-1 mice (Slc:ICR, 8 weeks, female) were infected intratracheally with 4.3-6.7 × 10^6^ CFUs of *S. pneumoniae.* For intratracheal infection, the vocal cords were visualized using an operating otoscope (Welch Allyn, NY, USA), and 40 µL of bacteria was placed onto the trachea using a plastic gel loading pipette tip. Mouse survival was monitored twice daily for 14 days. At 24 h after intratracheal infection, bronchoalveolar lavage fluid (BALF) was collected following perfusion with PBS.

For the sepsis model, CD-1 mice (Slc:ICR, 8 weeks, female) were infected intravenously with 3.3-6.5 × 10^5^ CFUs of *S. pneumoniae* via the tail vein. Mouse survival was monitored twice daily for 14 days. At 24 and 48 h after infection, blood aliquots were collected from mice following induction of general euthanasia. Brain, lung, and liver samples were collected following perfusion with PBS. Brain and lung whole tissues as well as the anterior segment of the liver were resected. Bacterial counts in the blood as well as organ homogenates were determined by separately plating serial dilutions, with organ counts corrected for differences in organ weight. Detection limits were 50 CFUs/organ and 50 CFUs/mL in blood.

The concentrations of TNF-α in BALF and plasma were determined using a Duoset^®^ ELISA Kit (R&D Systems, Minneapolis, MN, USA). Mice plasma was obtained by centrifuging the heparinized blood. All mouse experiments were conducted in accordance with animal protocols approved by the Animal Care and Use Committees at Osaka University Graduate School of Dentistry (28-002-0).

### Statistical analysis

Statistical analysis of *in vitro* and *in vivo* experiments was performed using a nonparametric analysis, Mann-Whitney *U* test, or Kruskal-Wallis test with Dunn’s multiple comparisons test. Mouse survival curves were compared using a log-rank test. *p* < 0.05 was considered to indicate a significant difference. The tests were carried out with Graph Pad Prism version 6.0h (GraphPad Software, Inc., San Diego, CA, USA).

## Results

### The *pfbA* gene is specific to *S. pneumoniae* among mitis group *Streptococcus*

We searched *pfbA*-homologues by tBLASTn and performed phylogenetic analysis (Fig. 1 and Supplementary Fig. 1). The *pfbA* gene homologues were identified in *S. pneumoniae, Streptococcus pseudopneumoniae*, and *Streptococcus merionis.* Although 16S rRNA sequences cannot distinguish mitis group species, the 16S rRNA of *Streptococcus* sp. strain W10853 showed 99.387% identity to that of *S. pseudopneumoniae.* Interestingly, *S. pneumoniae*-related species such as *Streptococcus mitis* and *Streptococcus oralis* did not contain the homologues, whereas *S. merionis* had a gene of which the query cover and identity were over 50%. *S. merionis* strain NCTC13788 (also known as WUE3771, DSM 19192, and CCUG 54871), isolated from the oropharynges of Mongolian jirds (*Meriones unguiculatus*), contained 16S rRNA that belongs in a cluster distinct from the mitis group (46). This result indicates that the *pfbA* gene is specific to *S. pneumoniae* and *S. pseudopneumoniae* in the mitis group.

**Figure 1.**
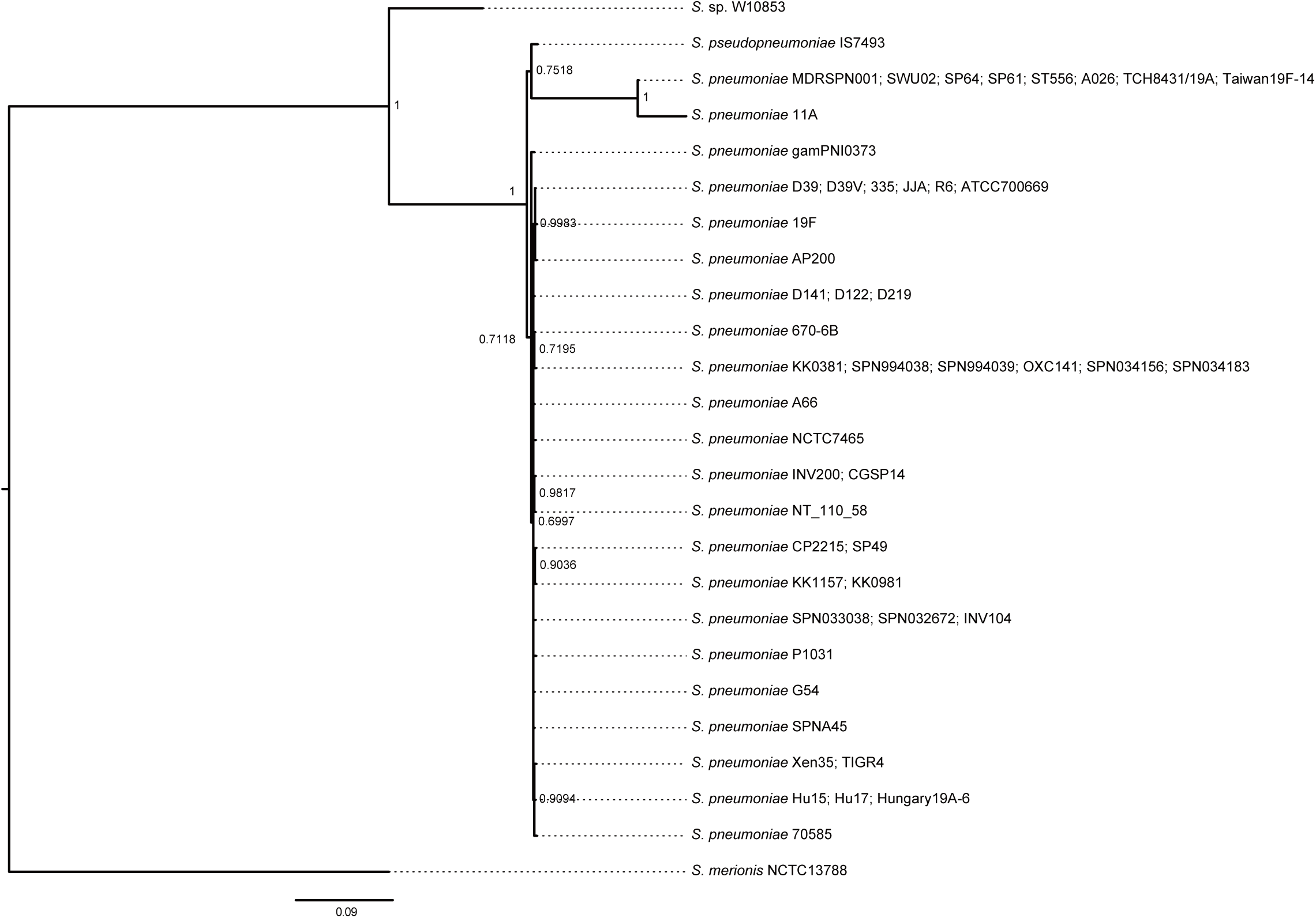
Bayesian phylogenetic analysis of the *pfbA* gene. The codon-based Bayesian phylogenetic relationship was calculated using the MrBayes program. Strains with identical sequences are listed on the same branch. The percentage of posterior probabilities is shown near the nodes. The scale bar indicates nucleotide substitutions per site.

### PfbA contributes to evasion of neutrophil killing

To investigate whether PfbA contributes to evasion of neutrophil killing, we determined pneumococcal survival rates after incubation with human neutrophils. After 3 h incubation, the TIGR4 Δ*pfbA* strain showed a significantly decreased bacterial survival rate. In addition, to clarify whether the observed effects were attributed to PfbA, we also performed the assay with rPfbA. In the presence of 100 nM rPfbA, TIGR4 Δ*pfbA* strain demonstrated a recovered survival rate nearly equal to that of the wild-type strain (Fig. 2A). In pneumococcal survival assays with neutrophil-like differentiated HL60 cells, TIGR4 strains showed similar results (Fig. 2B). We also performed the assay using the non-encapsulated strain R6 and human neutrophils. The R6 Δ*pfbA* strain showed significantly decreased survival rates as compared to the wild-type strain after incubation for 1, 2, and 3 h (Fig. 2C). As the R6 strain showed this phenotype at earlier time points than the TIGR4 strain, we performed pneumococcal survival assays using R6 strains with inhibitors (Fig. 2D). Neutrophil phagocytic killing of *S. pneumoniae* requires the serine proteases (47). Thus, we used a protein inhibitor cocktail as a positive control of a neutrophil killing inhibitor. While the R6 Δ*pfbA* strain showed significantly decreased survival rates at 1 h after incubation with human fresh neutrophils in the absence of inhibitors, treatment with an actin polymerization inhibitor, cytochalasin D, reduced the differences among the wild-type and Δ*pfbA* strains as well as the protein inhibitor cocktail. These results indicate that PfbA contributes to pneumococcal evasion of neutrophil phagocytosis.

**Figure 2.**
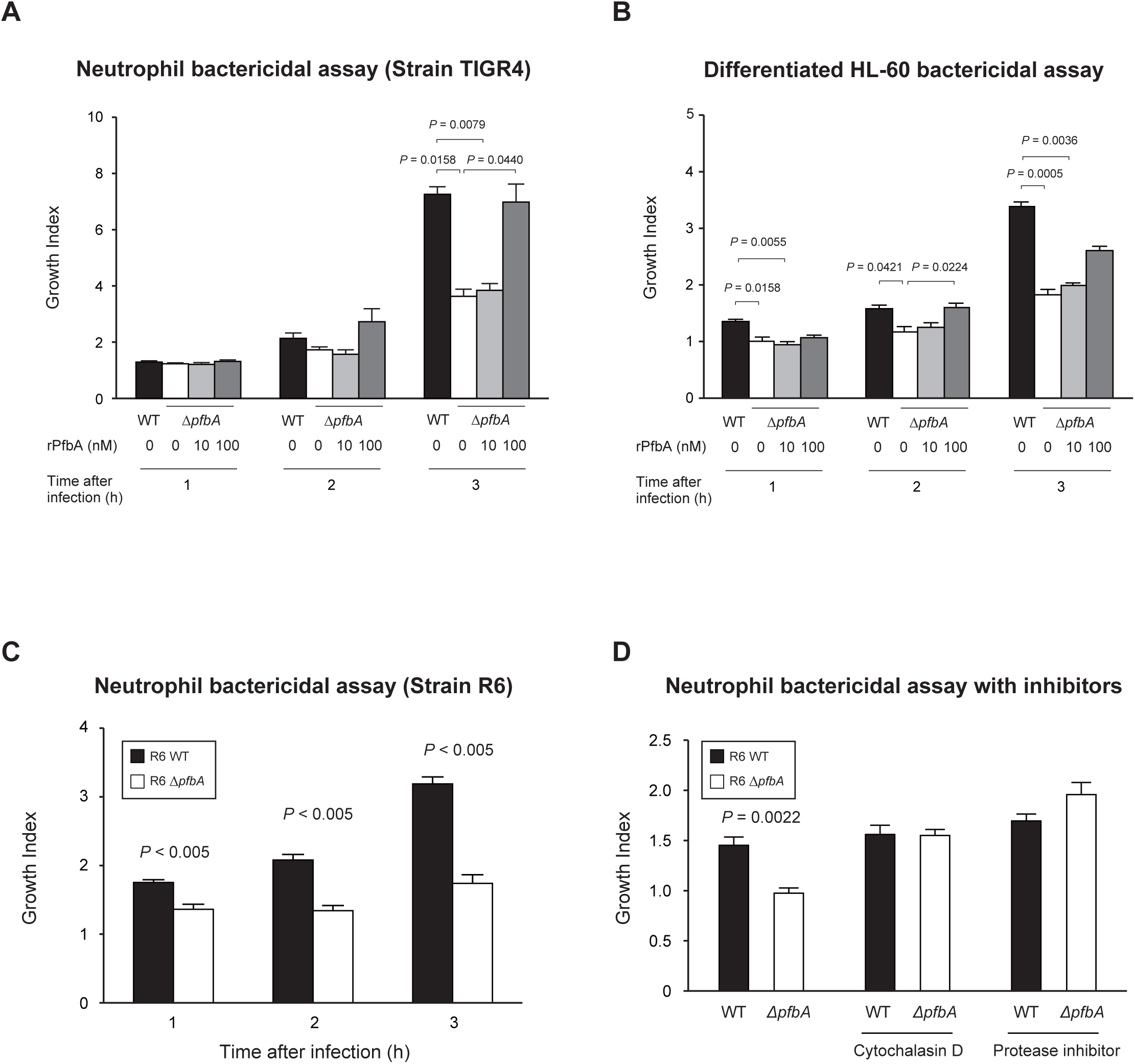
PfbA contributes to pneumococcal survival after incubation with neutrophils. **A.** Growth of TIGR4 strains incubated with human fresh neutrophils. **B.** Growth of TIGR4 strains incubated with neutrophil-like differentiated HL-60 cells. Bacterial cells were incubated with human neutrophils or differentiated HL-60 cells in the presence or absence of rPfbA for 1, 2, and 3 h at 37°C in a 5% CO_2_ atmosphere. Next, the mixture was serially diluted and plated on TS blood agar. Following incubation, the number of CFUs was determined. Growth index was calculated by dividing CFUs after incubation by CFUs of the original inoculum. **C.** Growth of R6 strains incubated with human fresh neutrophils. *S. pneumoniae* strains were added to human neutrophils without serum and gently mixed for 1, 2, or 3 h at 37°C. Next, the mixtures were serially diluted and plated on TS blood agar. After incubation, the number of CFUs was determined. **D.** Growth of R6 strains incubated with human fresh neutrophils in the presence of inhibitors. *S. pneumoniae* strains were added to human neutrophils with or without cytochalasin D, or protease inhibitor cocktail in the absence of serum, then gently mixed for 1 h at 37°C. The percent bacterial survival was calculated based on viable counts relative to the wild-type strain. These data are presented as the mean values of six samples, with S.E. values represented by vertical lines. Differences between several groups were analyzed using a Kruskal-Wallis test followed by Dunn’s multiple comparisons test (A, B). The Mann-Whitney’s U test was used to compare differences between two independent groups (C, D). Three experiments were performed, with data from a representative experiment is shown.

### PfbA inhibits neutrophil phagocytosis directly

We confirmed the anti-phagocytic activity of PfbA using flow cytometry and PfbA-coated fluorescent beads (Fig. 3A). The fluorescence intensity of neutrophils and monocytes incubated with PfbA-coated beads was substantially lower as compared with cells incubated with non- or BSA-coated beads. These results indicated that neutrophils and monocytes phagocytosed the non- and BSA-coated fluorescent beads, whereas the PfbA-coated fluorescent beads escaped phagocytosis by neutrophils and monocytes.

**Figure 3.**
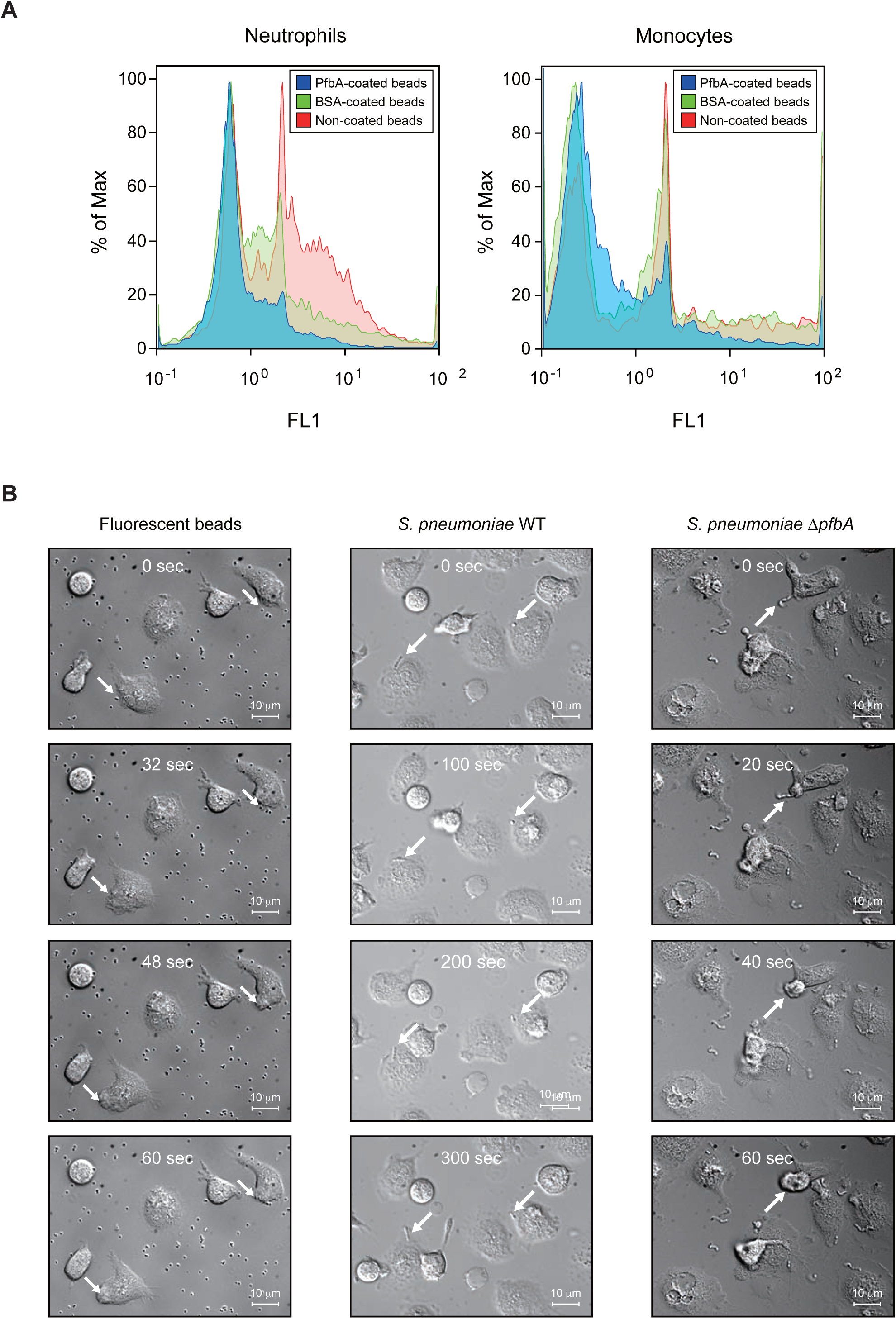
PfbA suppresses host cell phagocytosis. **A.** Uptake of fluorescent PfbA-coated beads by neutrophils and monocytes. Human neutrophils and monocytes were separately incubated with PfbA-, BSA-, or non-coated fluorescent beads for 1 h at 37°C. Phagocytic activities were analyzed using flow cytometry. Data are presented as histograms. The value shown for the percent of maximum was determined by dividing the number of cells in each bin by the number of cells in the bin that contained the largest number of cells. The bin is shown as a numerical range for the parameter on the X-axis. **B.** Time-lapse analysis of the interaction between *S. pneumoniae* and neutrophils. *S. pneumoniae* wild-type and Δ*pfbA* strains were incubated with neutrophils. The elapsed times from contact with neutrophils are shown in the upper part of the figures. Arrows indicate when *S. pneumoniae* cells contacted neutrophils. Arrowheads indicate *S. pneumoniae* engulfed by a neutrophil phagosome.

We performed real-time observations for time-lapse analysis of the interaction between *S. pneumoniae* and neutrophils (Fig. 3B). *S. pneumoniae* strain R6 wild-type and Δ*pfbA* strains were separately incubated with fresh human neutrophils in RPMI 1640 medium. After coming into contact with neutrophils, the Δ*pfbA* strain was phagocytosed within 1 min, whereas the wild-type strain was not phagocytosed after more than 5 min. Time-lapse analysis also showed the Δ*pfbA* strain engulfed by neutrophil phagosomes. These results suggest that PfbA can directly inhibit phagocytosis.

### PfbA works as a TLR2 ligand and may inhibit phagocytosis through TLR2

Some lectins of pathogens work as ligand for TLR2 and TLR4 (48). We previously reported that PfbA can interact with glycolipid and glycoprotein fractions of red blood cells, several monosaccharides, D-sucrose, and D-raffinose (26, 27). Hence, to determine whether PfbA works as a TLR ligand, we performed a SEAP assay using HEK-293 cells stably transfected with either TLR2 or TLR4, NF-κB, and SEAP (Fig. 4A). Pam3CSK4 and Zymozan were used as positive controls for the TLR2 ligand, while LPS was used for TLR4. The SEAP assay indicated that pasteurized *S. pneumoniae* TIGR4 wild-type cells activated NF-κB via TLR2, whereas Δ*pfbA* cells did not stimulate cells expressing either TLR2 or TLR4. Pasteurized rPfbA also activated NF-κB dose-dependently through TLR2, but not TLR4. In addition, in the presence of pasteurized rPfbA, Δ*pfbA* cells activated the cells expressing TLR2. Thus, PfbA is responsible for pneumococcal NF-κB activation through TLR2.

**Figure 4.**
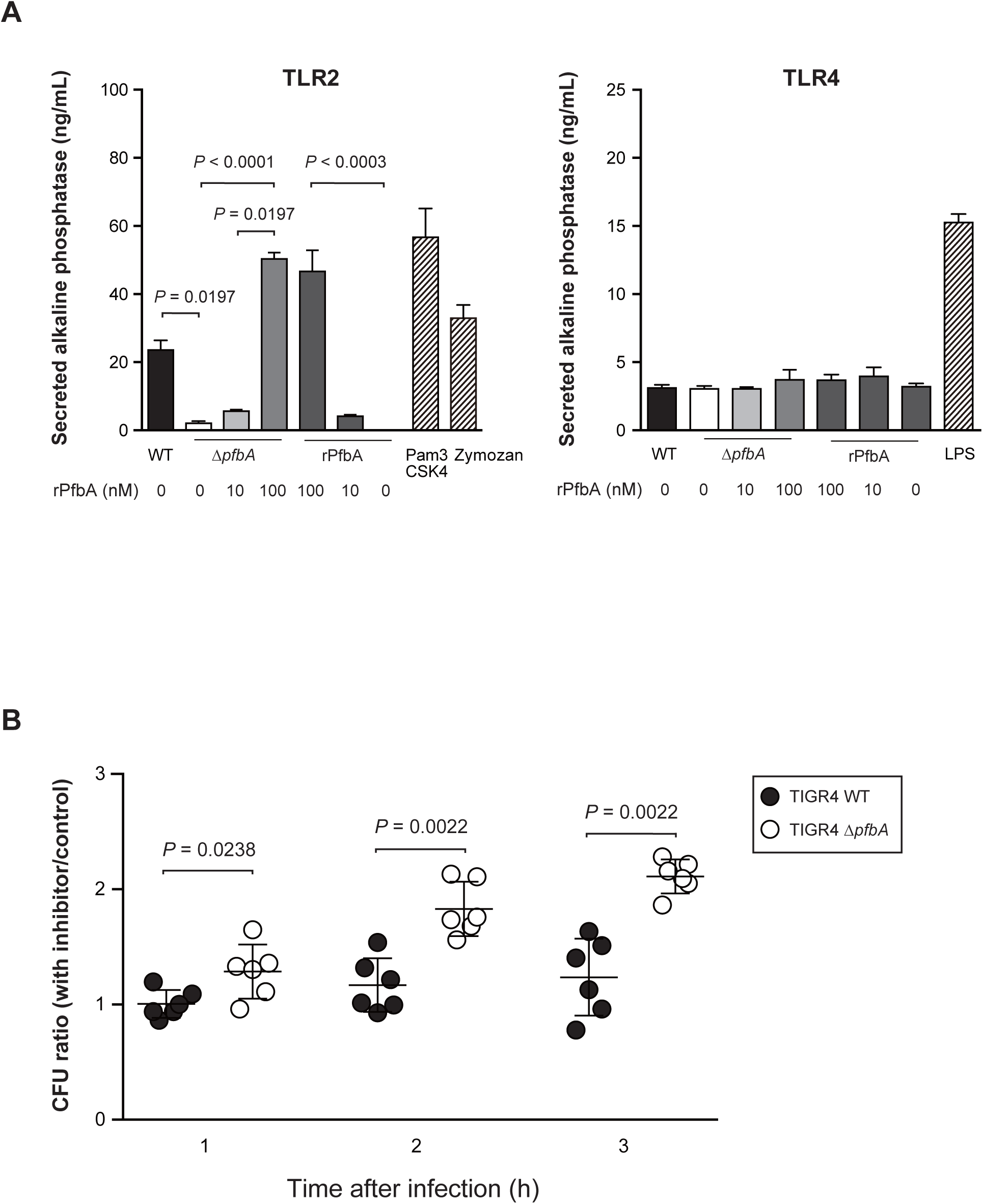
PfbA activates NF-κB via TLR2, and TLR2/4 inhibitor enhances Δ*pfbA* strain survival. **A.** Secreted alkaline phosphatase (SEAP) porter assay using TLR2/NF-κB/ SEAPorter or TLR4/MD-2/CD14/NF-κB SEAPorter HEK293 cell lines. The cells were plated in 24-well plates at 5 × 10^5^ cells/well. After 24 h, cells were stimulated with various amount of rPfbA, pasteurized *S. pneumoniae* (∼5 × 10^6^ CFU), 1 µg/mL Pam3CSK4, 10 µg/mL Zymozan, or 25 ng/mL LPS for 24 h. SEAP was analyzed using the SEAPorter Assay Kit. Data are presented as the mean of six wells. SE values are represented by vertical lines. Differences in pneumococcal infection group and rPfbA addition group were analyzed using a Kruskal-Wallis test followed by Dunn’s multiple comparisons test, respectively. **B.** TLR2/4 inhibitor peptide enhances survival of the TIGR4 Δ*pfbA* strain incubated with human neutrophils. *S. pneumoniae* TIGR4 wild type strain or Δ*pfbA* strain bacteria were incubated with human neutrophils in the presence of TLR2/4 inhibitor peptide or control peptide. After 1, 2, and 3 h, the mixture was serially diluted and plated on TS blood agar. Following incubation, the number of CFUs was determined. The CFU ratio was calculated by dividing CFUs in the presence of inhibitor peptide by CFUs in the presence of control peptide. Data are presented as the mean of six wells. S.E. values are represented by vertical lines. Differences between groups were analyzed using Mann-Whitney’s U test.

Next, to determine whether TLR signaling suppresses survival of pneumococci incubated with neutrophils, we performed a neutrophil survival assay using a TIRAP inhibitor peptide (Fig. 4B). Data are presented as the ratio calculated by dividing CFUs in the presence of inhibitor peptide by CFUs in the presence of control peptide. TIRAP is an adaptor protein involved in MyD88-dependent TLR2 and TLR4 signaling pathways. Since the TIRAP inhibitor peptide blocks the interaction between TIRAP and TLRs, the peptide works as a TLR2 and TLR4 inhibitor. The inhibitor peptide treatment increased survival rates of the Δ*pfbA* strain, but did not affect wild-type survival rates. These results indicate that PfbA contributes to the evasion of neutrophil phagocytosis, and TIRAP inhibitor treatment did not change survival rates of pneumococci incubated with neutrophils. On the other hand, the *S. pneumoniae* Δ*pfbA* strain is more easily phagocytosed by neutrophils as compared to the wild-type strain, and this phenotype is abolished by TIRAP inhibitor.

Stimulation of the human monocytic cell line THP1 by a TLR ligand, LPS, induces miR-146a/b expression in an NF-κB-dependent fashion, and this induction inhibits innate immune responses (49). In addition, pneumococcal infection of human macrophages induces expression of several microRNAs, including miR-146a, in a TLR-2-dependent manner, which prevents excessive inflammation (50). We performed microRNA array analysis using neutrophil like-differentiated HL60 cells, *S. pneumoniae* strains and rPfbA (Supplementary Fig. 2, Accession number: GSE128341). We compared rPfbA-treated and non-treated cells, wild type and Δ*pfbA*-infected cells, and Δ*pfbA* with and without rPfbA-infected cells. The analysis revealed only one microRNA, hsa-miR-1281, that was commonly downregulated by 2-fold or greater in the presence of PfbA as compared to in its absence (Supplementary Fig. 2, magenta circle). On the other hand, there were no commonly upregulated miRNAs, including miR-146a/b. In addition, the expression of eight microRNAs was commonly changed in wild-type infection and Δ*pfbA* infection with rPfbA as compared to infection with Δ*pfbA* only. Five micro RNAs (hsa-miR-4674, hsa-miR-3613-3p, hsa-miR-4668-5p, hsa-miR-3197, and hsa-miR-6802-5p) were upregulated, while three (hsa-miR-3935, hsa-miR-1281, and hsa-miR-3613-5p) were downregulated. However, the role of these miRNAs in infectious process remains unclear.

### PfbA deficiency reduces pneumococcal burden in BALF but does not alter host survival rate in a mouse pneumonia model

To investigate the role of PfbA in pneumococcal pathogenesis, we infected mice with *S. pneumoniae* strains intratracheally and compared bacterial CFUs and TNF-α levels in BALF from mice 24 h after infection. There were no differences observed in survival time between mice infected with wild type and Δ*pfbA* strains (Fig. 5A). However, recovered CFUs of wild-type bacteria were significantly greater than those of Δ*pfbA* strains in mouse BALF. In addition, the level of TNF-α in BALF was almost the same in wild type and Δ*pfbA* infection (Fig. 5B).

**Figure 5.**
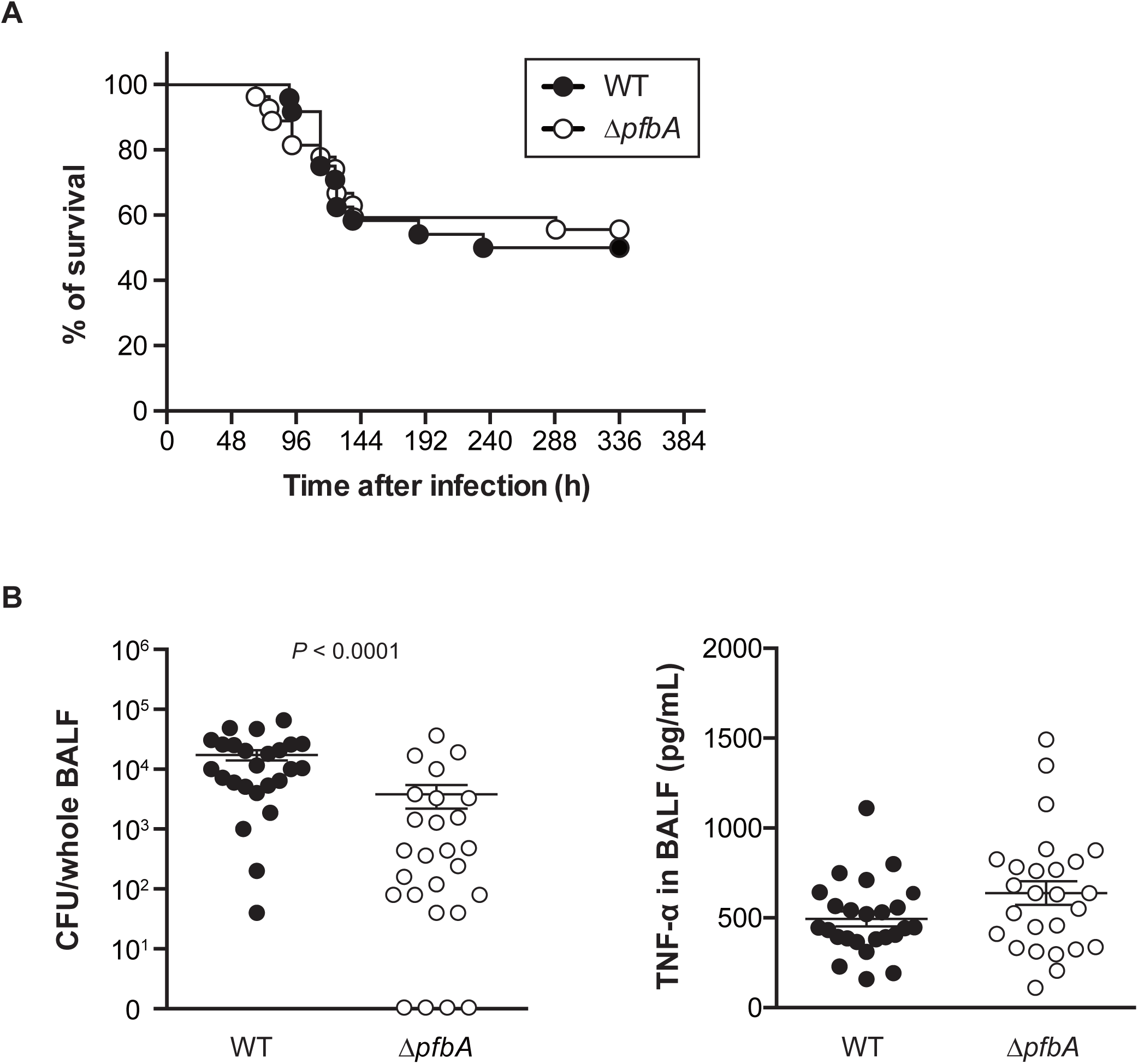
In a mouse pneumonia model, deficiency of *pfbA* decreases pneumococcal burden in the lung but does not affect host mortality. **A.** CD-1 mice were infected intratracheally with the *S. pneumoniae* TIGR4 wild-type or Δ*pfbA* strain (3-18 × 10^6^ CFUs). Mice survival was recorded for 14 days. The differences between groups were analyzed using a log-rank test. **B.** Bacterial CFUs and TNF-α in BALF collected from CD-1 mice after intratracheal infection with *S. pneumoniae*. CD-1 mice were infected intratracheally with the *S. pneumoniae* TIGR4 wild type or Δ*pfbA* strain (4-7 × 10^6^ CFUs). BALF was collected at 24 h after pneumococcal infection, and bacterial CFUs and TNF-α levels in the BALF were determined. S.E. values are represented by vertical lines. Statistical differences between groups were analyzed using Mann-Whitney’s U test. The data obtained from three independent experiments were pooled.

### PfbA deficiency increases pneumococcal pathogenicity in a mouse sepsis model

We also investigated the role of PfbA in mice following intravenous infection as a model of sepsis. In the infection model, the Δ*pfbA* strain showed significantly higher levels of virulence as compared to the wild-type strain (Fig. 6A). Furthermore, we compared the TNF-α levels in plasma and examined the bacterial burden in blood, brain, lung, and liver samples obtained at 24 and 48 h after intravenous infection (Fig. 6B, 6C and Supplementary Fig. 3). At 24 h after infection, TNF-α ELISA findings showed a significantly greater level in the plasma of *pfbA* mutant strain-infected mice as compared to the wild-type strain-infected mice. The numbers of CFUs of both the wild-type and *pfbA* mutant strains in the blood and brain samples were comparable. On the other hand, in the lung and liver samples, the *pfbA* mutant strain-infected mice showed slightly but significantly reduced numbers of CFUs as compared with the wild-type strain-infected mice. At 48 h after infection, there were no significant differences in TNF-α level and bacterial burden in each organ between the wild-type- and *pfbA* mutant strain-infected mice (Supplementary Fig. 3). Bacteria were not detected in the blood of two of the wild-type strain-infected mice and five of the *pfbA* mutant strain-infected mice. Meanwhile, three of the wild-type strain-infected mice yielded more than 10^6^ CFUs/mL, while seven of the wild-type strain-infected mice did. The *pfbA* mutant strain infection caused a polarized bacterial burden in the host at 48 h after infection as compared to wild type infection.

**Figure 6.**
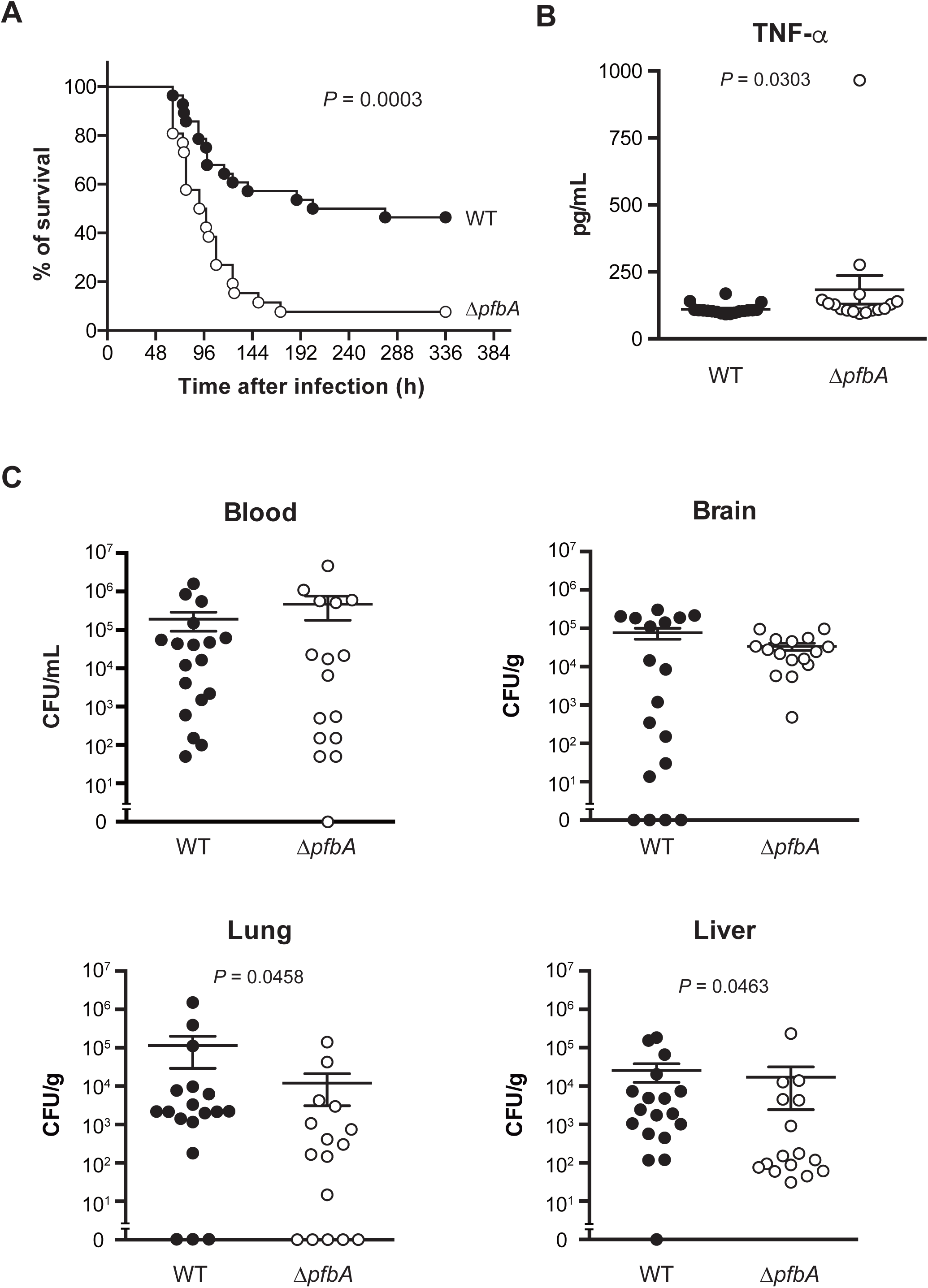
In a mouse sepsis model, the deficiency of *pfbA* increases the virulence and TNF-α level in blood but decreases the bacterial burden in the lung and liver. CD-1 mice were infected intravenously with the *S. pneumoniae* TIGR4 wild type or Δ*pfbA* strain (3-6 × 10^6^ CFUs). **A.** Mouse survival was monitored for 14 days. Statistical differences between groups were analyzed using a log-rank test. **B.** CD-1 mice were infected intravenously with the *S. pneumoniae* TIGR4 wild type or Δ*pfbA* strain (6-9 × 10^6^ CFUs). Plasma samples were collected from these mice at 24 h after infection. Values are presented as the mean of 16 or 18 samples. Vertical lines represent the mean ± S.E. Statistical differences between groups were analyzed using Mann-Whitney’s U test. **C.** The bacterial burden in the blood, brain, lung, and liver were assessed after 24 h of infection. S.E. values are represented by vertical lines. All mice were perfused with PBS after blood collection, organ samples were collected. Statistical differences between groups were analyzed using Mann-Whitney’s U test. The mouse survival data were obtained from three independent experiments, and the TNF-α level and bacterial burden values obtained from two independent experiments were pooled.

## Discussion

In the present study, we found that *pfbA* is a pneumococcal-specific gene that contributes to evasion of neutrophil phagocytosis. We determined that PfbA can activate NF-κB through TLR2. TIRAP inhibition increased the survival rate of Δ*pfbA* strain incubated with neutrophils, while this inhibition did not affect a wild-type strain survival. In a mouse model with lung infection, the bacterial burden of the Δ*pfbA* strain was significantly reduced as compared with that of wild-type strain, but the TNF-α level was comparable between the strains. Overall, there was no significant difference in the survival rates of mice infected with the wild-type *S. pneumoniae* strain- and those infected with the Δ*pfbA* strain. Furthermore, in a mouse model with blood infection, the Δ*pfbA* strain showed a significantly higher TNF-α level than the wild-type strain. These results suggest that PfbA may suppress the host innate immune response by acting as an anti-phagocytic factor interacting with TLR2.

Prior studies have shown that *S. pneumoniae* under selective pressure can adapt to the environment by importing genes from other related streptococci, such as those in the mitis group (51-54). Although *S. mitis* and *S. oralis* are oral commensal bacteria, these species contain various pneumococcal virulence factor homologues. Some mitis group strains harbor several choline-binding proteins including autolysins, pneumolysin, sialidases, and others (11, 55, 56). In this study, we found that *pfbA* homologues were absent among mitis group strains without *S. pneumoniae* for which whole genome sequences were available, whereas the *pfbA* gene is highly conserved among pneumococcal strains. Interestingly, a streptococcal species with clear evolutionary separation from the mitis group, *S. merionis*, contained a *pfbA* orthologue. This result indicates that *pfbA* is a pneumococcal-specific gene and that ancestral *S. pneumoniae* likely obtained the gene by horizontal gene transfer from non-mitis group streptococcal species.

Although lipoproteins are major TLR2 ligands as well as peptidoglycans in *S. pneumoniae* (19), we found that rPfbA can activate NF-κB solely in HEK293 cells expressing TLR2, but not those expressing TLR4. Since *E. coli* does not have the capacity to glycosylate proteins (57), rPfbA-mediated TLR2 activation would be independent of pneumococcal glycosylation. Plant and pathogen lectins can induce NF-κB activation through binding to TLR2 *N*-glycans, while a classical ligand such as Pam3CSK4 can activate NF-κB glycan-independently (48). TLR2 has four *N*-glycans whose structures still remain unknown, and the *N*-glycans are critical for the lectins to induce TLR2-mediated activation (48). PfbA binds to various carbohydrates via the groove residues in the β-helix (26, 27). There is a possibility that PfbA induces TLR2 signaling by binding to TLR2 *N*-glycans.

Human macrophages challenged with *S. pneumoniae* induce a negative feedback loop, preventing excessive inflammation via miR-146a and potentially other miRNAs on the TLR2-MyD88 axis (50). On the other hand, pneumococcal endopeptidase O enhances macrophage phagocytosis in a TLR2- and miR-155-dependent manner (58). Furthermore, miR-9 is induced by TLR agonists and functions in feedback control of the NF-κB-dependent responses in human monocytes and neutrophils (59). These studies indicate that host phagocytes are regulated by a complex combination of pattern recognition receptor signaling and miRNA induction. We predicted that PfbA suppresses phagocytosis via the induction of miRNAs in a TLR2 dependent fashion. However, an miRNA array showed that the levels of the involved miRNAs were not changed over 2-fold in the presence or absence of PfbA. One possible hypothesis is that PfbA induces different miRNA responses from classical TLR ligands via glycan-dependent recognition. Although PfbA can downregulate miR-1281 in differentiated HL-60 cells, the role of miR-1281 in phagocytes remains unclear. Further comprehensive studies are required to investigate the role of miRNAs in host innate immunity.

Unexpectedly, our mouse pneumonia and sepsis models indicated that *pfbA* deficiency reduces pneumococcal survival in the host, but does not decrease or increases host mortality. We previously reported that PfbA works as an adhesin and invasin of host epithelial cells (22). The reduction of bacterial burden in host organs can be explained by the synergy of adhesive and anti-phagocytic abilities. On the other hand, the *S. pneumoniae* Δ*pfbA* strain showed equivalent or greater induction of inflammatory cytokines as compared with the wild-type strain. Generally, a deficiency of TLR ligands would suppress inflammatory responses. However, a deficiency of PfbA would cause more efficient bacterial uptake by phagocytes and promote inflammatory responses. In addition, there is a possibility that the negative feedback loop induced by PfbA is lost and causes excess inflammation. High mortality does not mean bacterial success, as host death leads to the limitation of bacterial reproduction. PfbA may be beneficial for pneumococcal species by increasing the bacterial reproductive number through suppression of host cell phagocytosis and host mortality. PfbA showed high specificity for and conservation in *S. pneumoniae* species. The assumed negative feedback loop may not be as significant in non-pathogenic mitis group *Streptococcus*.

In single toxin-induced infectious diseases such as diphtheria and tetanus, highly safe and protective vaccines are established. On the other hand, in multiple factor-induced diseases such as those caused by *S. pneumoniae, S. pyogenes*, and so on, there are either no approved vaccines or existing vaccines still need optimization. Our study indicates that PfbA is a pneumococcal specific cell surface protein, which contributes to evasion from phagocytosis. Therefore, PfbA would not be suitable as a vaccine antigen, since the protein suppresses pneumococcal virulence in a mouse sepsis model. Further investigation of the intricate balance between host immunity and pathogenesis is required to establish the basis for drug and vaccine design.

## Acknowledgements

This study was supported by the Japanese Society for the Promotion of Science (JSPS), KAKENHI [grant numbers 26861546, 15H05012, 16H05847, 16K15787, 17H05103, 17K11666, and 18K19643], the SECOM Science and Technology Foundation, Takeda Science Foundation, GSK Japan Research Grant, Asahi Glass Foundation, Kurata Memorial Hitachi Science and Technology Foundation, Kobayashi International Scholarship Foundation, and the Naito Foundation.

## Author contributions

M.Y. and S.K. designed the study. M.Y. performed bioinformatics analyses. M.Y., Y.H., M.T., and M.O. performed the experiments. M.Y., T.S., M.N., Y.T., and S.K. contributed to the setup of the experiments. M.Y. wrote the manuscript. Y.H., M.T., M.O., T.S., M.N., Y.T., and S.K. contributed to the writing of the manuscript.

