## Supplementary information for "*Streptococcus pneumoniae* evades host cell phagocytosis and limits host mortality through its cell wall anchoring protein PfbA"

Running title: PfbA inhibits phagocytosis and limits host responses

Masaya Yamaguchi[a,#], Yujiro Hirose[a], Moe Takemura[a, b], Masayuki Ono[a, c], Tomoko Sumitomo[a], Masanobu Nakata[a], Yutaka Terao[d], Shigetada Kawabata[a]

[a]Department of Oral and Molecular Microbiology, Osaka University Graduate School of Dentistry, Osaka, Japan

[b]Department of Oral and Maxillofacial Surgery II, Osaka University Graduate School of Dentistry, Osaka, Japan

[c]Department of Fixed Prosthodontics, Osaka University Graduate School of Dentistry, Osaka, Japan.

[d]Division of Microbiology and Infectious Diseases, Niigata University Graduate School

of Medical and Dental Sciences, Niigata, Japan

**Supplementary Figure 1. Maximum likelihood phylogenetic analyses of the *pfbA* gene**

The codon-based maximum likelihood phylogenetic relationship was calculated using the RAxML program. Strains with identical sequences are listed on the same branch. The bootstrap values are shown near the nodes. The scale bar indicates nucleotide substitutions per site.

**Supplementary Figure 2. microRNA array analysis of differentiated HL60 cells**

miRNA array analysis was performed using the Affymetrix GeneChip® miRNA 4.0 array. Bacterial cells and/or rPfbA were incubated with differentiated HL-60 cells for 1 h at 37°C in a 5% $CO_2$ atmosphere. Total RNA including micro RNA was purified with an miRNeasy kit. X and Y axes represent expression values after normalization. The red dashed line represents 2-fold change. Magenta circle means commonly downregulated miRNA by 2-fold or greater in the presence of PfbA as compared to in its absence.

**Supplementary Figure 3. Mice were intravenously infected with *S. pneumoniae* TIGR4 wild-type or *ΔpfbA* strains.**

**A.** Plasma samples were collected from intravenously infected mice at 48 hours after infection. Values are presented as the mean of 16 or 18 samples. Vertical lines represent the mean ± S.E. Statistical differences between groups were analyzed using Mann-Whitney’s U test. **B.** The bacterial burden in the blood, brain, lung, and liver were assessed after 48 h of infection. S.E. values are represented by vertical lines. All mice were perfused with PBS after blood collection, organ samples were collected. Statistical differences between groups were analyzed using Mann-Whitney's U test.

**Supplementary Table 1. Primers used in this study**

| Primers | Sequence (5' to 3') |
|---|---|
| For knockout plasmid construction | |
| T4pfbAKOuF | CCGCGGGAATTCGATgtgtcttgttctagttttcaattca |
| T4pfbAKOuR | tattcaaatatatcccatcagaacctccaattttttact |
| T4pfbAKOaF | ttggaggttctgatgggatatatttgaatacatacgaaca |
| T4pfbAKOaR | atttatctttaccattcaattttttataattttttaat |
| T4pfbAKOdF | ttataaaaaaattgaatggtaaagataaataaaatttgtt |
| T4pfbAKOdR | GAATTCACTAGTGATtcaaacatcaatgactagaatactt |
| T4pfbAKOvF | gtcattgatgtttgaATCACTAGTGAATTCGCGGCCGCCT |
| T4pfbAKOvR | actagaacaagacacATCGAATTCCCGCGGCCGCCATGGC |
| For double-crossover recombination | |
| pfbAKOuMaxF | gtgtcttgttctagttttcaattca |
| pfbAKOdMaxR | tcaaacatcaatgactagaatactt |
| For mutation confirmation | |
| T4pfbAKOupup | tagcatttagaatccttactagac |
| T4pfbAKOdndn | ccccaatacgttcaatgtcagttg |

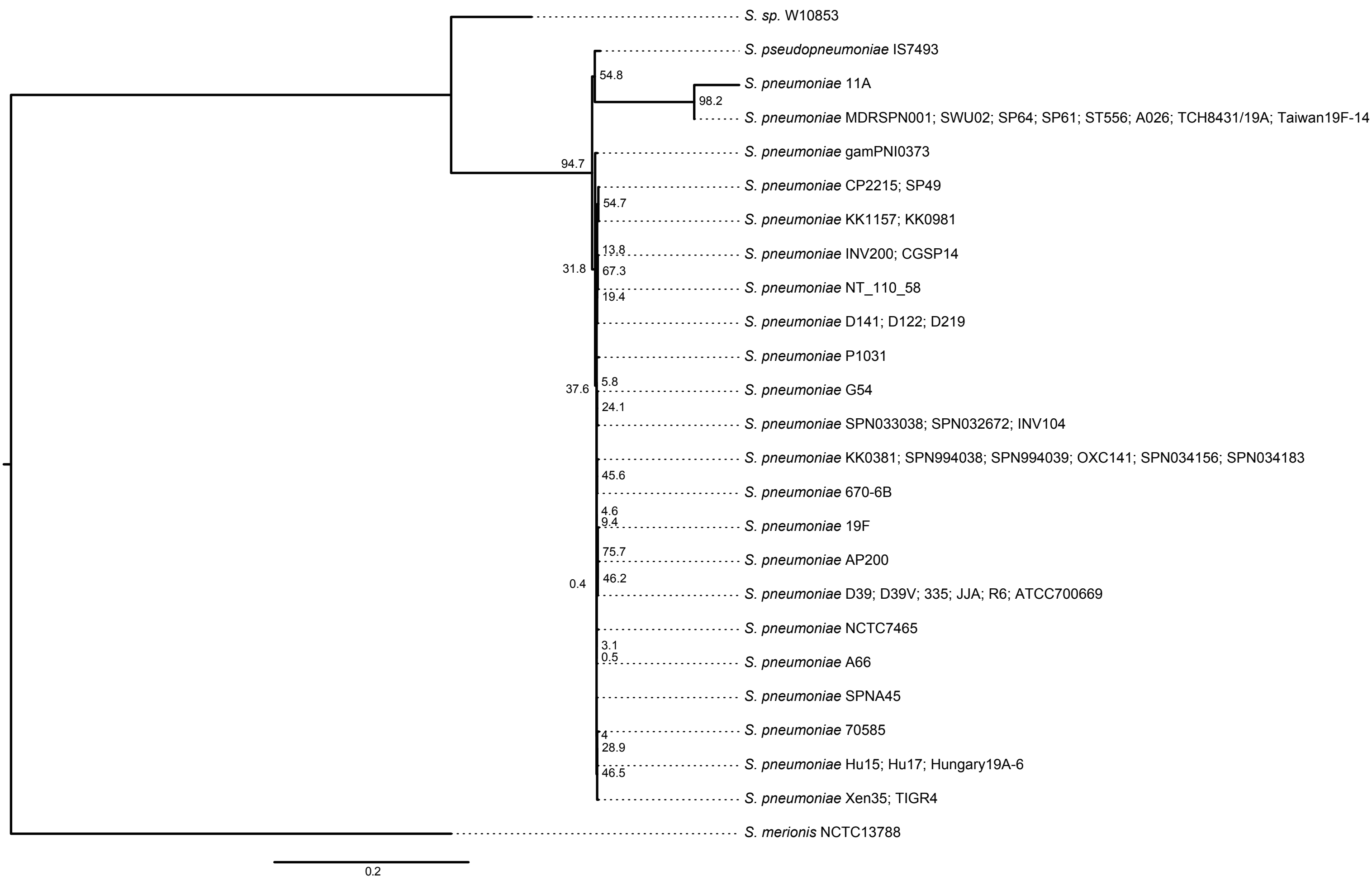


Supplementary Figure 1. Yamaguchi *et al.*

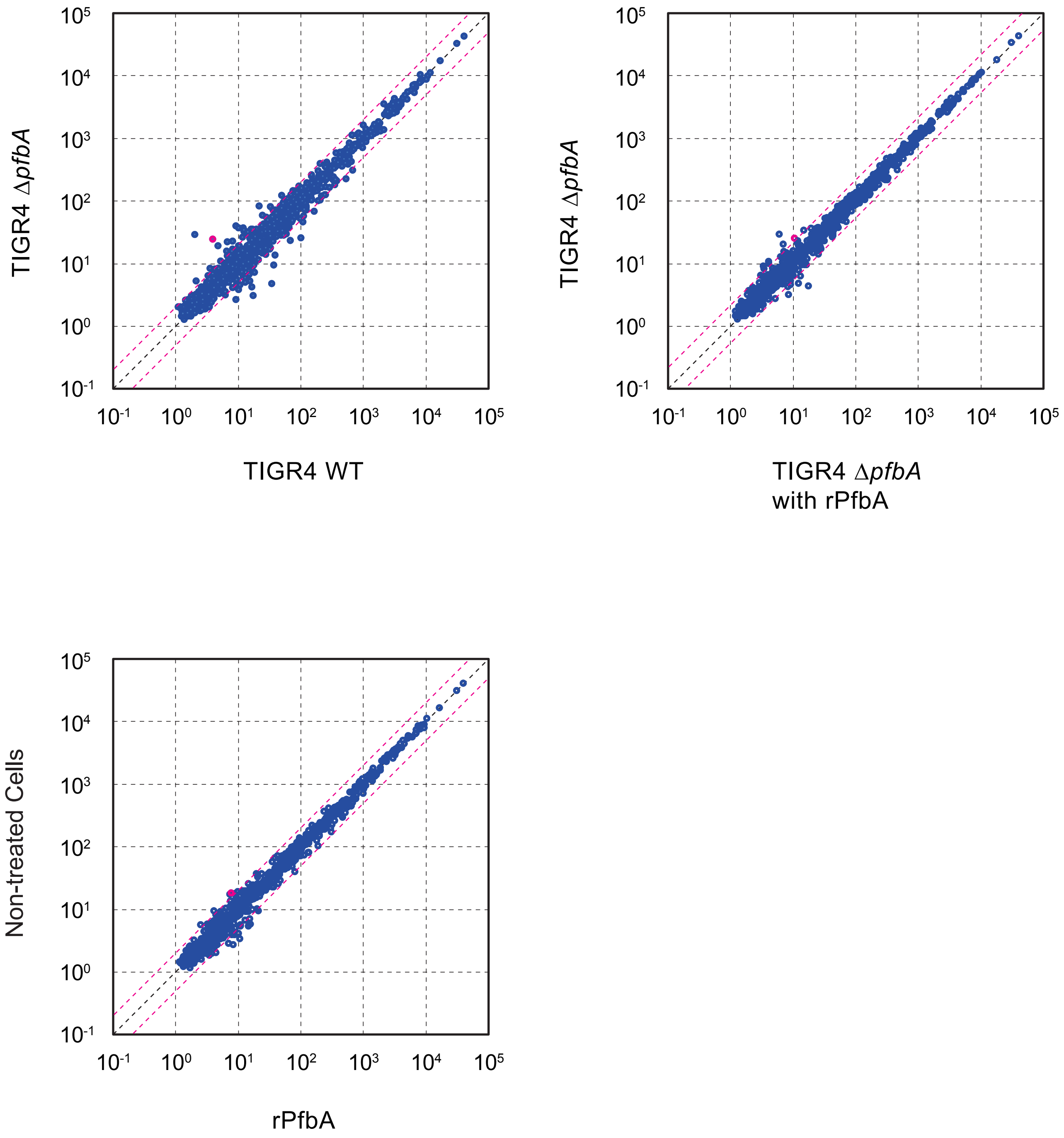


Supplementary Figure 2. Yamaguchi *et al.*

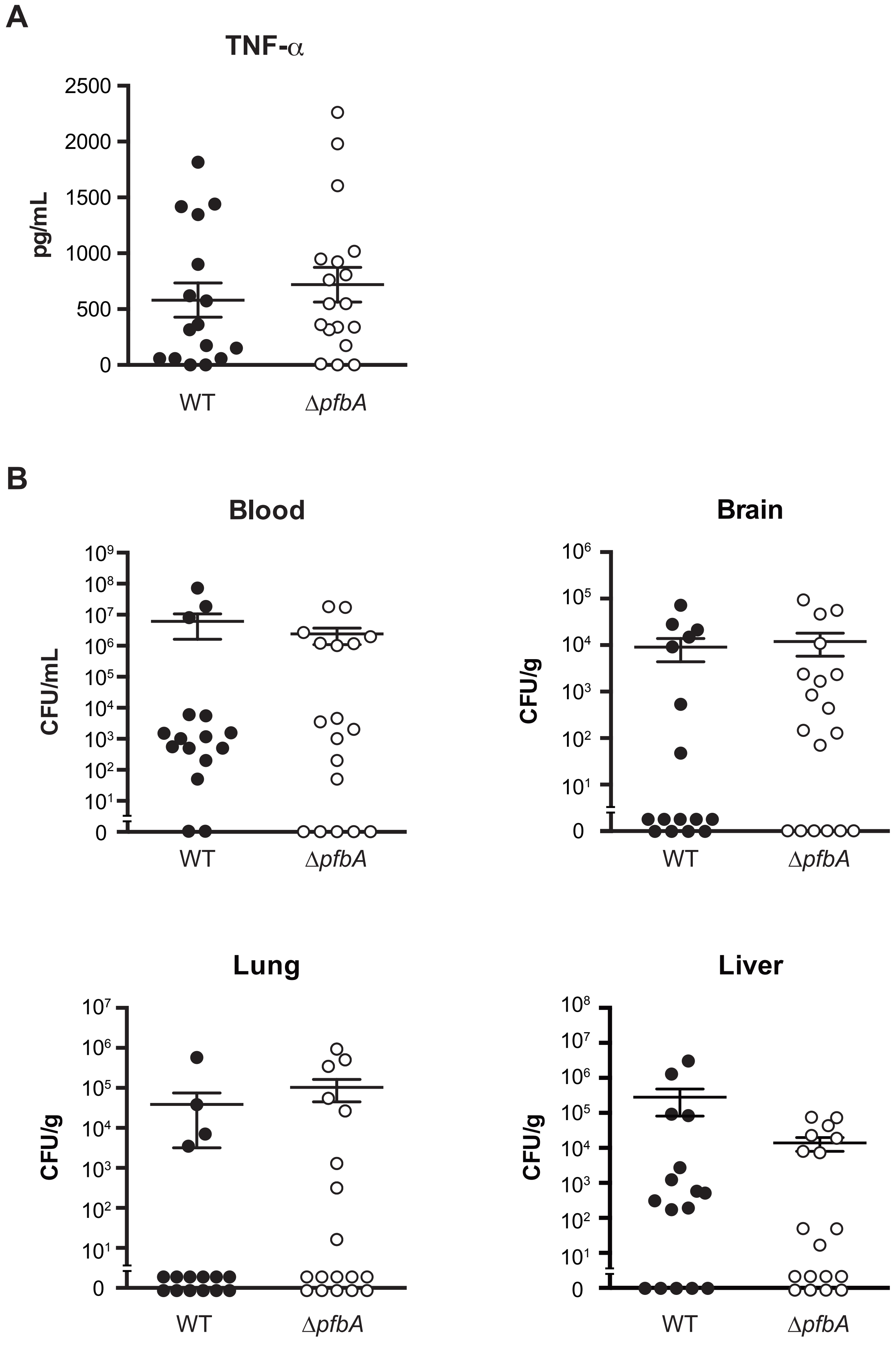


Supplementary Figure 3. Yamaguchi *et al.*
